# Identification of ubiquitin Ser57 kinases regulating the oxidative stress response in yeast

**DOI:** 10.1101/2020.06.20.162883

**Authors:** Nathaniel L. Hepowit, Kevin N. Pereira, Jessica M. Tumolo, Walter J. Chazin, Jason A. MacGurn

**Author notes:** Address correspondence to Jason A. MacGurn.

## Abstract

Ubiquitination regulates many different cellular processes, including protein quality control, membrane trafficking, and stress responses. The diversity of ubiquitin functions in the cell is partly due to its ability to form chains with distinct linkages that can alter the fate of substrate proteins in unique ways. The complexity of the ubiquitin code is further enhanced by post-translational modifications on ubiquitin itself, the biological functions of which are not well understood. Here, we present genetic and biochemical evidence that serine 57 (Ser57) phosphorylation of ubiquitin functions in stress responses in *Saccharomyces cerevisiae*, including the oxidative stress response. We also identify and characterize the first known Ser57 ubiquitin kinases in yeast and human cells, and we report that two Ser57 ubiquitin kinases regulate the oxidative stress response in yeast. These studies implicate ubiquitin phosphorylation at the Ser57 position as an important modifier of ubiquitin function, particularly in response to proteotoxic stress.

## INTRODUCTION

Ubiquitin is a post-translational modifier that regulates diverse cellular processes in eukaryotic cells. The broad utility of ubiquitin as a regulatory modification is due to the high degree of complexity associated with ubiquitin polymers, which are added to substrate proteins by the activity of ubiquitin conjugation machinery (E1-E2-E3 cascades) and removed from substrates by deubiquitylases (DUBs). Ubiquitin can be conjugated recursively at any of seven internal lysines or the N-terminus to generate polymers with distinct topological features *(1–3).* Complexity is further enhanced by the formation of mixed and branched polymers *(4–6)* and by post-translational modifications that can occur on ubiquitin itself *(1)*. For example, PINK1-mediated phosphorylation of ubiquitin at the Ser65 position plays an important role in mitophagy by regulating parkin-mediated ubiquitination of mitochondrial membrane proteins *(7–12)*. Phosphorylation of ubiquitin at the Ser57 has also been reported *(13–15)*, but the kinases that produce this modification and its regulatory significance remain unknown.

Many proteotoxic stresses activate ubiquitin networks to promote protein quality control and protect the cell from damage associated with systemic protein misfolding. Oxidative stress is highly damaging to the cell, triggering deployment and re-distribution of existing ubiquitin pools and induction of ubiquitin biosynthesis to promote survival by activating a repertoire of ubiquitin-mediated responses *(16)*. During oxidative stress, many proteins become damaged and misfolded, resulting in a global increase in K48-linked ubiquitin conjugation that targets substrates for clearance by proteasome-mediated degradation *(17, 18)*. Oxidative stress also triggers translation arrest *(19)*, resulting in K63-linked polyubiquitylation on ribosomes to stabilize the 80S complex and the formation of polysomes *(20).* Furthermore, oxidative damage of DNA activates signaling and repair processes that are tightly regulated by K63 ubiquitylation and deubiquitylation activities *(21–25).* Thus, the oxidative stress response relies heavily on ubiquitin networks to regulate many critical biological processes.

Although ubiquitin networks play a critical role in the cellular response to proteotoxic stress, precisely how these networks are tuned to enhance protein quality control and other protective functions remains unclear. Given that Ser65 phosphorylation of ubiquitin regulates the clearance of damaged mitochondria *(7–12)*, we hypothesized that phosphorylation at other positions may regulate ubiquitin function, particularly in conditions that promote protein damage and misfolding. Since it is the most abundant phosphorylated form *(14)*, we examined the biological functions of Ser57 phosphorylated ubiquitin in yeast, aiming to identify and characterize the molecular events and signaling processes that regulate its production.

## RESULTS AND DISCUSSION

To probe potential biological functions of Ser57 ubiquitin phosphorylation in yeast, we analyzed the growth of yeast strains expressing exclusively wildtype, Ser57Ala (phosphorylation resistant, or S57A) or Ser57Asp (phosphomimetic, or S57D) ubiquitin under various stress conditions. Expression of S57A or S57D ubiquitin did not affect growth of yeast at ambient (26°C) temperatures (**FIG 1A**), consistent with previous reports *(13, 15, 26).* However, we noticed that expression of S57D ubiquitin enhanced both long-term and acute tolerance of thermal stress (**FIG 1A-C**). We also observed that yeast expressing S57D ubiquitin were hypersensitive to hydroxyurea, while cells expressing S57A ubiquitin exhibited resistance to tunicamycin (**FIG 1, supplement 1**).

**Fig. 1.**
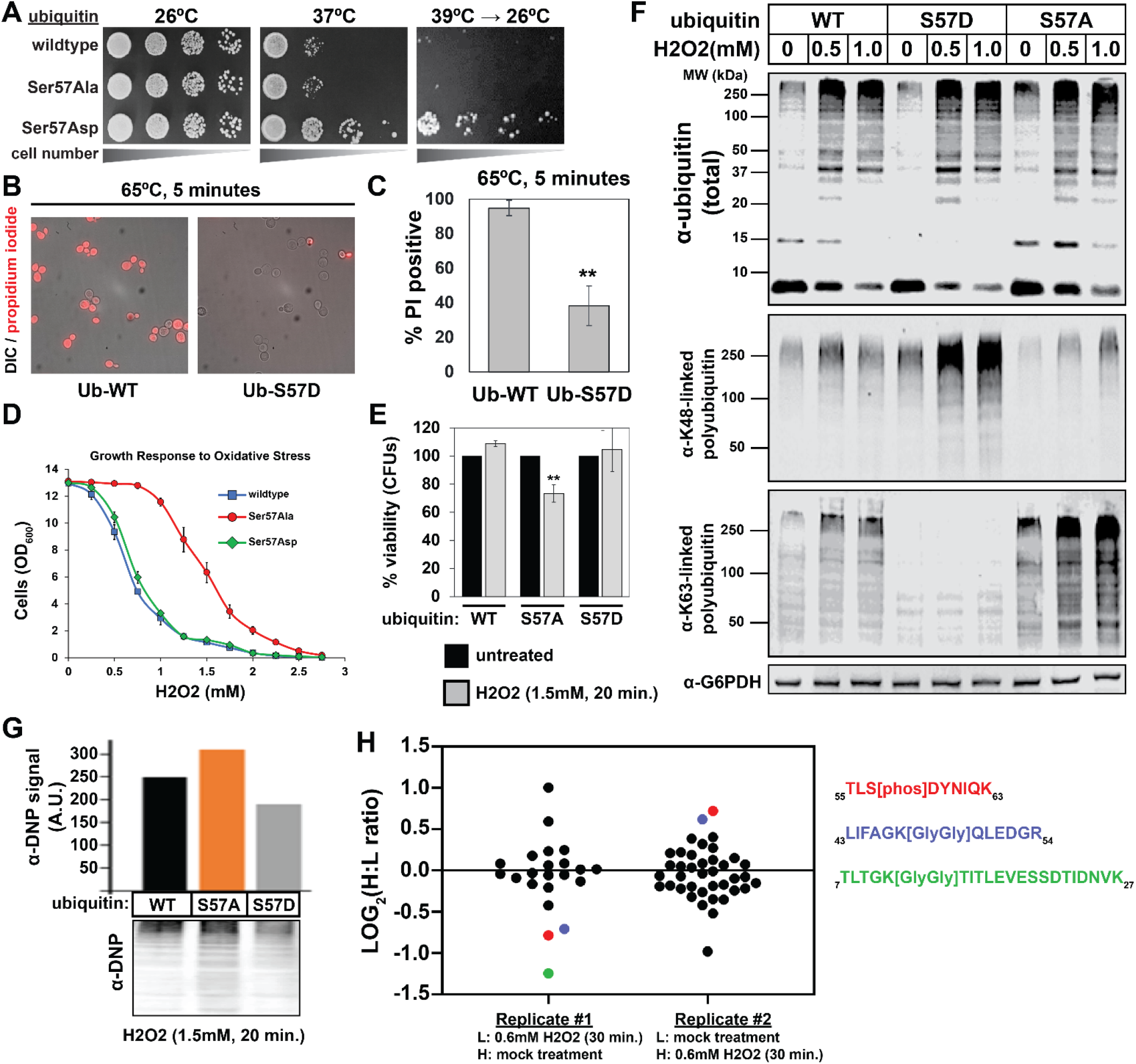
Ser57 phosphorylation of ubiquitin is important for the yeast proteotoxic stress response. **(A)** Serial dilution of yeast cells on YPD agar plate and incubated at 26°C, 37°C, or 39°C for 18 hours and shifted back to 26°C for recovery. **(B)** Fluorescence microscopy and **(C)** quantification of cells stained with propidium iodide (PI) after incubation at 65°C for 5 min. Dead cells are stained with PI. **(D)** OD_600_ of cells cultured in YPD with different concentrations of H_2_O_2_ for 48 hours (starting OD_600_ = 0.025). **(E)** CFU count of cells treated with 1.5 mM H_2_O_2_ for 20 minutes. Untreated control was taken prior to H_2_O_2_ treatement. **(F)** Western blot with anti-ubiquitin, anti-K48 ubiquitin and anti-K63 ubiquitin antibodies of lysates from cells treated with different concentrations of H_2_O_2_ for 30 minutes. **(G)** Anti-dinitrophenyl (DNP) western blot of oxidized (carbonylated) proteins derivatized with 2,4-dinitrophenylhydrazine. **(H)** SILAC-MS of affinity purified 3XFLAG-ubiquitin from yeast cells treated with 0.6 mM H_2_O_2_ for 30 minutes. Cells used in A-G are SUB280-derived strains exclusively expressing WT, S57A or S57D ubiquitin.

We next analyzed the role of Ser57 phosphorylation in the oxidative stress response. Wildtype yeast cells arrest growth in response to oxidative stress and activate responses that help the cells cope with the proteotoxic and DNA damaging effects of oxidation *(20, 27–29)*. Interestingly, while yeast cells expressing wildtype or S57D ubiquitin arrested growth in response to moderate oxidative stress (>1mM H_2_O_2_), cells expressing S57A ubiquitin were deficient in this response and only arrested growth in response to more severe oxidative stress (>2mM H_2_O_2_) (**FIG 1D**). The failed growth arrest observed for cells expressing S57A ubiquitin correlated with decreased viability (**FIG 1E**).

Since oxidative stress induces the production of both K48- and K63-linked ubiquitin conjugates *(18, 20, 30, 31)* we decided to test if expression of S57A and S57D ubiquitin alters ubiquitin conjugation patterns in response to oxidative stress. We found that 30 minute exposure of yeast cells to hydrogen peroxide (0.5mM and 1.0mM) resulted in increased abundance of K48- and K63-linked ubiquitin polymers (**FIG 1F**), consistent with a previous report *(20)*. Compared to cells expressing wildtype ubiquitin, we found that cells expressing S57D ubiquitin exhibited increased abundance of K48-linked polymers and decreased abundance of K63-linked polymers (**FIG 1F**). In contrast, cells expressing S57A ubiquitin exhibited increased abundance of K63-linked polyubiquitin chains and decreased abundance of K48-linked polyubiquitin chains (**FIG 1F**). These results suggest that Ser57 phosphorylation may contribute to the regulation of ubiquitin conjugation and polymer linkage type used during oxidative stress. We hypothesized that the increased production of K48-linked polyubiquitin in cells expressing S57D ubiquitin might promote the clearance of damaged and misfolded proteins that accumulate during oxidative stress. To test this, we reacted cellular proteins with DNP hydrazine (which covalently modifies carbonylated proteins that occur with oxidative stress) followed by SDS-PAGE and immunodetection of DNP adducts. We found that cells expressing S57A ubiquitin exhibited increased accumulation of oxidized proteins, while cells expressing S57D ubiquitin exhibited decreased accumulation of oxidized proteins (**FIG 1G**). Importantly, we also found that oxidative stress induced an approximately 2-fold increase in phosphorylation of ubiquitin at the Ser57 position (**FIG 1H** and **FIG 1, supplement 2**). Taken together, these results indicate that Ser57 phosphorylation induced during oxidative stress may play a role in the clearance of damaged and misfolded proteins, yet the kinases that mediate this phosphorylation reaction remain unknown.

To identify candidate ubiquitin kinases, we screened for Ser57 phosphorylation activity by co-expressing ubiquitin and yeast kinases in *E. coli* and immunoblotting lysates using an antibody specific for Ser57 phosphorylated ubiquitin. Initially, we focused on candidate kinases for which mutants exhibit phenotypes corresponding to those observed for cells expressing S57A or S57D ubiquitin. We found that co-expression of ubiquitin with the kinase Vhs1 resulted in immunodetection of Ser57 phosphorylated ubiquitin (**FIG 2A**). Vhs1 is part of the yeast family of Snf1-related kinases *(32)*, and additional screening revealed three other kinases in this family that phosphorylated ubiquitin at the Ser57 position (**FIG 2, supplement 1**): Sks1 (which is 43% identical to Vhs1), Gin4 and Kcc4. A previous study reported consensus phosphorylation motifs for Vhs1, Gin4 and Kcc4 and all bear similarity to the amino acid sequence surrounding Ser57 in ubiquitin (**FIG 2, supplement 2**) *(33)*.

**Fig. 2.**
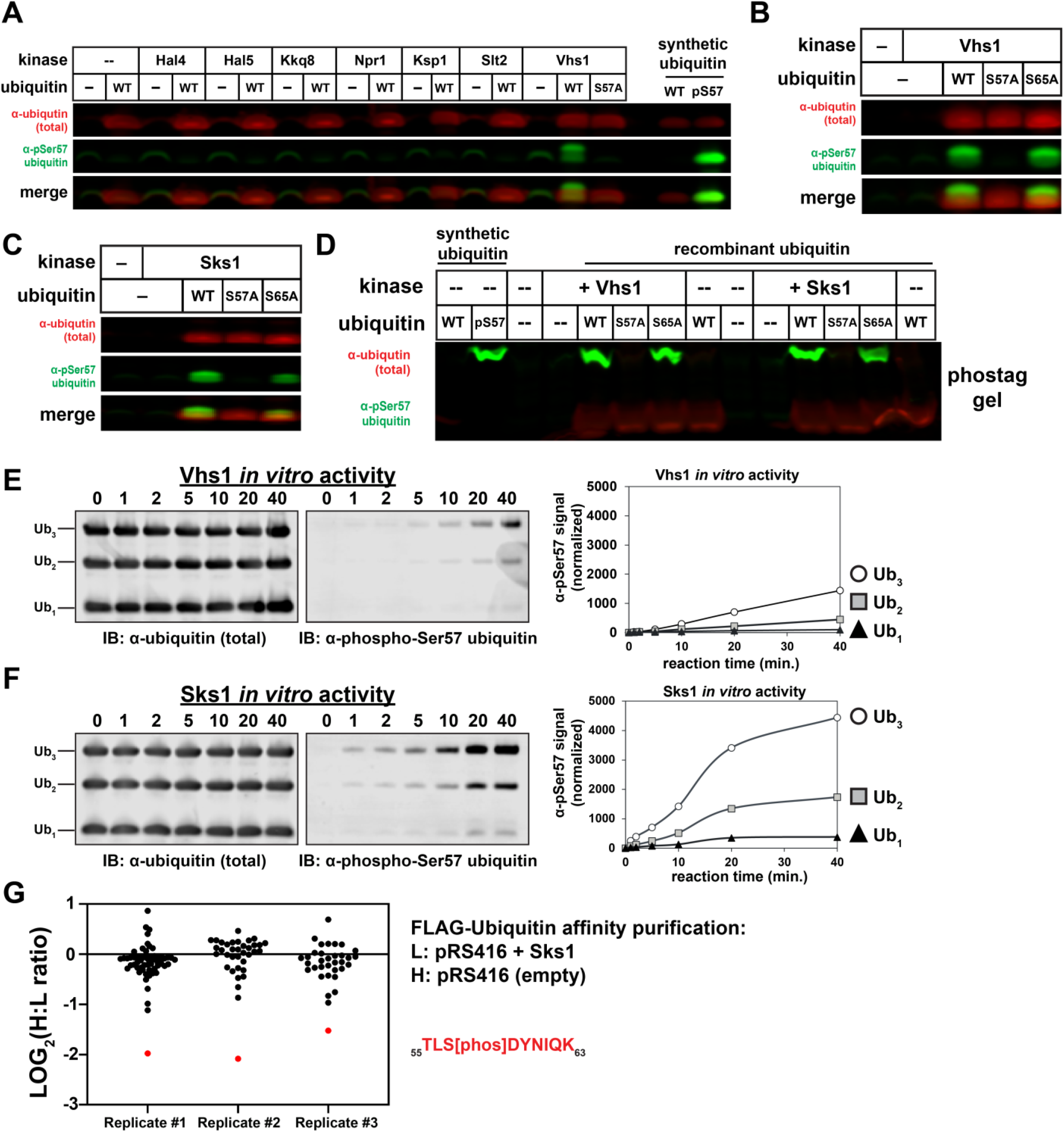
Identification and characterization of yeast ubiquitin Ser57 kinases. **(A)** Anti-phospho-Ser57 western blot of *E. coli* (Rosetta 2) lysates co-expressing ubiquitin and select members of the Snf1 family of kinases. **(B-C)** Ubiquitin variants were co-expressed with Vhs1 (B) and Sks1 (C) in *E. coli.* **(D)** Phos-tag gel separation of unmodified and Ser57-phosphorylated ubiquitin in the presence of Vhs1 or Sks1. **(E-F)** *In vitro* reconstitution of ubiquitin Ser57 phosphorylation using purified recombinant Vhs1 (E) and Sks1 (F). **(G)** SILAC-MS of affinity purified 3XFLAG-ubiquitin from yeast cells with or without Sks1 overexpression. Black and red dots indicate resolved ubiquitin peptides and Ser57-phosphopeptides, respectively.

We further characterized the activity of Vhs1 and Sks1. Point mutagenesis of ubiquitin revealed that Vhs1 and Sks1 activity was dependent on Ser57, but not Ser65 (**FIG 2B-C**). Analysis of Vhs1 and Sks1 activity using Phos-tag acrylamide gels (**FIG 2D**) and mass spectrometry (**FIG 2, supplements 3–4**) confirmed production of Ser57 phosphorylated ubiquitin. Using purified recombinant Vhs1 and Sks1, we reconstituted kinase activity toward Ser57 of ubiquitin and found that both kinases exhibited a preference for polymers (tri-ubiquitin > di-ubiquitin > mono-ubiquitin). The *in vitro* activity of Sks1 was noticeably greater than that of Vhs1 (**FIG 2E-F**). Analysis of linkage specificity in the phosphorylation reaction revealed that Vhs1 is active toward linear (M1-linked), K29-, and K48-linked tetramers, while Sks1 is active toward linear, K48, and K63-linked polymers (**FIG 2, supplement 5**). Together, these results show that Vhs1 and Sks1 have linkage-specific activities *in vitro* that may underlie subtle differences in their biological functions. To test if the observed *in vitro* activity correlated with *in vivo* activity, we over-expressed Sks1 in yeast cells and used SILAC to quantify changes to ubiquitin modifications. Importantly, we observed that Sks1 overexpression increased ubiquitin phosphorylation at the Ser57 position (**FIG 2G** and **FIG 2, supplement 6**). Taken together our data indicate that Vhs1 and Sks1 are bona fide ubiquitin kinases that specifically phosphorylate the Ser57 position.

To explore potential biological functions, we tested if deletion or overexpression of these Ser57 ubiquitin kinases phenocopies the expression of S57A or S57D ubiquitin, respectively (**FIG 1A-D** and **FIG 1, supplement 1**). While we did not observe any heat tolerance phenotypes in yeast cells lacking Ser57 ubiquitin kinases (**FIG 3, supplement 1**), we did observe the following phenotypes which correlated with Ser57 phenotypes: **(i)** Vhs1 overexpression conferred slight resistance to canavanine (**FIG 3, supplement 2**), reminiscent of canavanine resistance conferred by S57D expression *(15)*; **(ii)** *Δsks1Δvhs1* double mutant cells exhibited tunicamycin resistance (**FIG 3, supplement 3**) reminiscent of yeast cells expressing S57A ubiquitin (**FIG 1, supplement 1**); **(iii)** deletion of either *SKS1* or *VHS1* resulted in hydroxyurea resistance phenotypes (**FIG 3, supplement 4**), consistent with the hydroxyurea sensitivity phenotype observed in yeast cells expressing S57D ubiquitin (**FIG 1, supplement 1**); and **(iv)** *Δsks1Δvhs1* double mutant cells failed to arrest growth in response to oxidative stress (**FIG 3A**), similar to yeast cells expressing S57A ubiquitin (**FIG 1D**). Notably, this latter phenotype was suppressed by expression of S57D (but not WT or S57A) ubiquitin (**FIG 3B**), indicating that the growth arrest phenotype in *Δsks1Δvhs1* cells is linked to a defect in ubiquitin phosphorylation at the Ser57 position. Taken together, these phenotypic correlations suggest that Vhs1 and Sks1 function in the same stress response pathways and processes as Ser57 phosphorylation of ubiquitin.

**Fig. 3.**
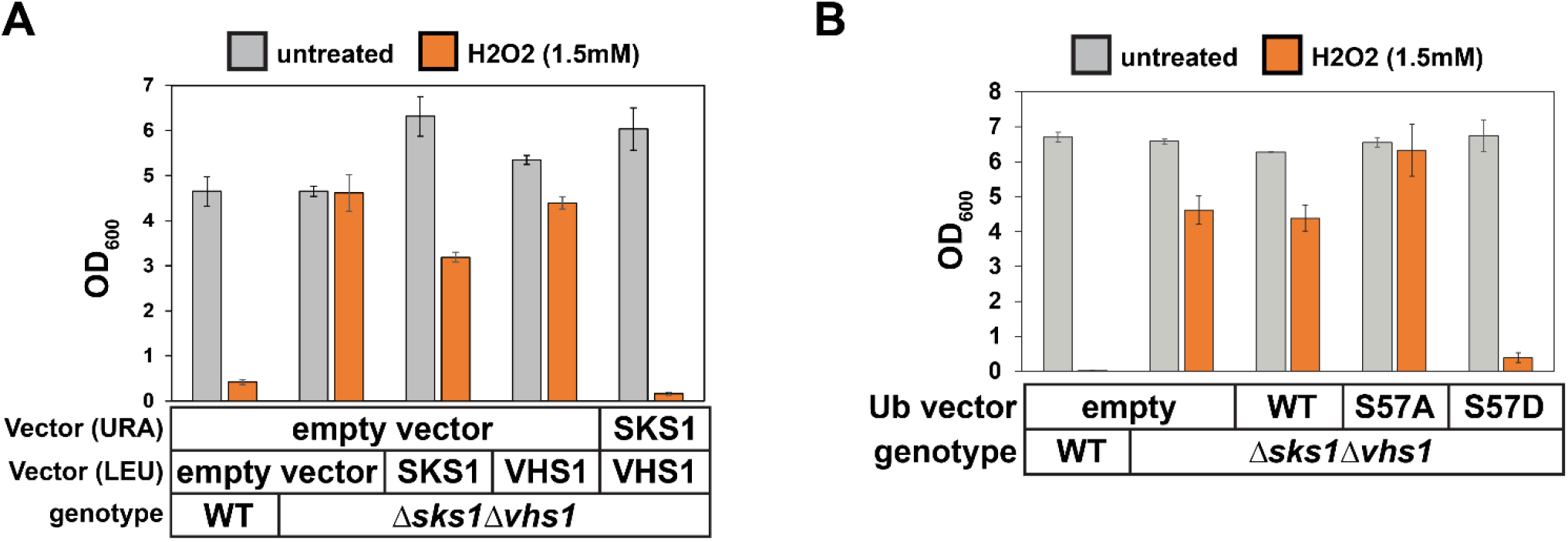
Vhs1 and Sks1 regulate the response to oxidative stress. **(A)** Analysis of the growth response to oxidative stress in WT and *Δsks1Δvhs1* double mutant cells, with indicated complementation expression vectors. OD_600_ of cells in dropout SM media in the absence or presence of 1.5 mM H_2_O_2_ after incubation at 24 h or 48 h, respectively. Starting OD_600_ of 0.025 in quadruplicates. **(B)** Analysis of the growth response to oxidative stress in WT and *Δsks1Δvhs1* double mutant cells, with indicated ubiquitin expression vectors. OD_600_ of cells in dropout SM media in the absence or presence of 1.5 mM H_2_O_2_ after incubation at 24 h or 48 h, respectively. Starting OD_600_ of 0.025 in quadruplicates.

The family of Snf1-related kinases in yeast share homology with the human AMPK-related kinases *(32)*. To test if human AMPK-related kinases exhibit activity toward ubiquitin, we performed *in vitro* kinase assays and found that a subset of this family phosphorylated ubiquitin at the Ser57 position specifically on tetramers (**FIG 4A**). This activity was apparent in the MARK kinases (MARK1-4) as well as related kinases QIK and SIK. Mass spectrometry analysis confirmed production of Ser57 phosphorylated ubiquitin by MARK2 *in vitro* (**FIG 4B**). Further analysis of MARK2 activity toward linkage-specific ubiquitin tetramers revealed a preference for linear, K11-, K29-, and K63-linked tetramers (**FIG 4C**). It is noteworthy that other candidate human ubiquitin kinases (initially identified by commercial screening services) did not exhibit Ser57 ubiquitin kinase activity *in vitro* (**FIG 4, supplement 1**). Given that Ser57 ubiquitin phosphorylation activity was detected within yeast and human Snf1-related kinases (**FIG 4, supplement 2**), we propose that this is an evolutionarily conserved function for a subset of kinases within this family. The data presented here indicate that Ser57 phosphorylated ubiquitin and the kinases that produce it play an important role in the oxidative stress response. Based on the diverse phenotypes associated with gain (and loss) of Ser57 phosphorylated ubiquitin, combined with the identification of multiple related ubiquitin Ser57 kinases in yeast and humans, we predict that Ser57 phosphorylation of ubiquitin and the kinases that catalyze this modification play important regulatory roles in stress responses and other cellular processes.

**Fig. 4.**
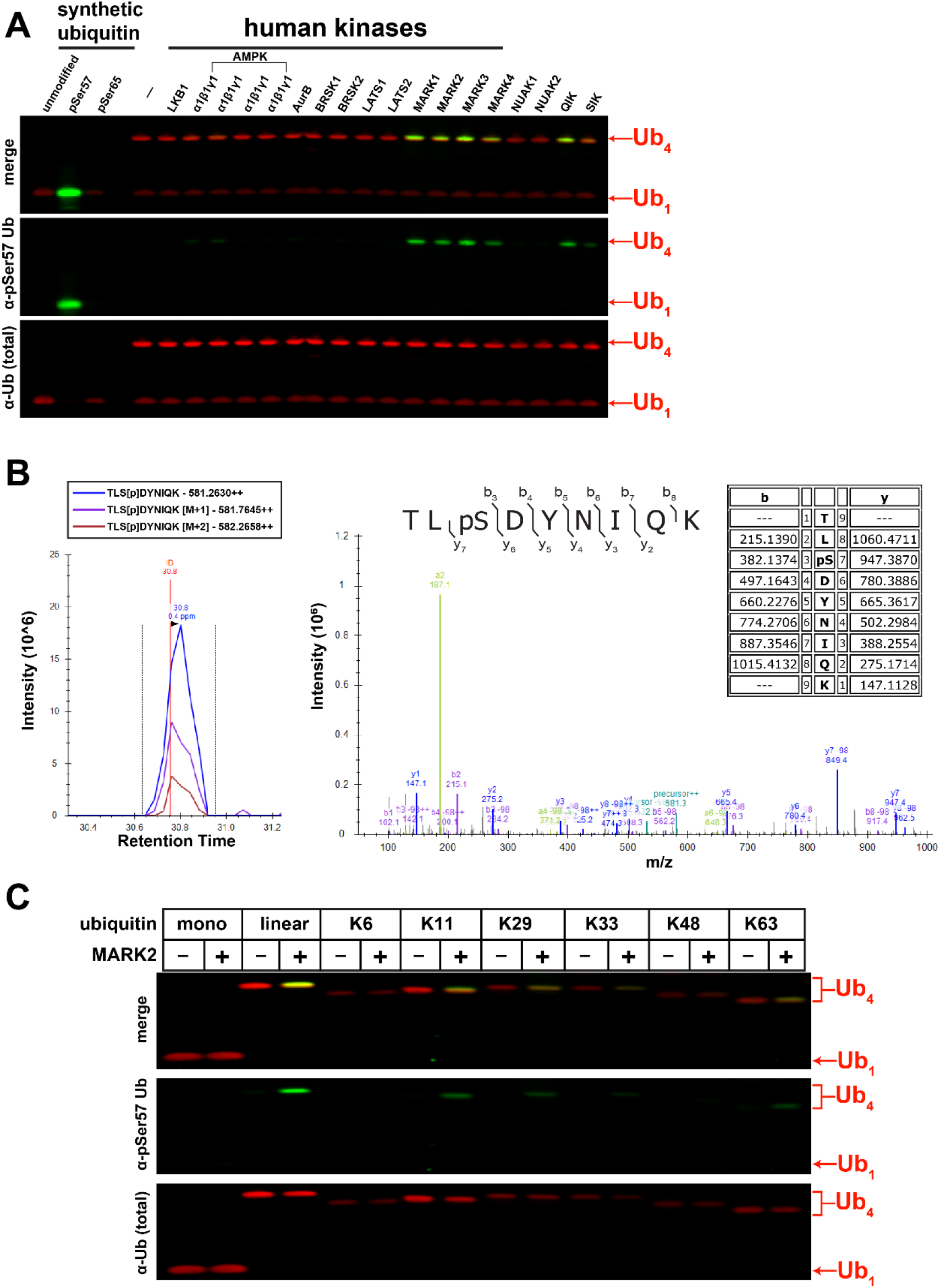
Identification and characterization of human ubiquitin Ser57 kinases. **(A)** *In vitro* activity assay of select human kinases of the AMPK family using mono-ubiquitin and M1-linked tetra-ubiquitin as substrates. **(B)** Mapping the phosphorylation sites on ubiquitin in the presence of MARK2 using mass spectrometry. **(C)** Determination of MARK2 substrate preference by analyzing activity toward tetra-ubiquitin chains with different linkages.

## MATERIALS AND METHODS

### Cell strains and culture

All *Saccharomyces cerevisiae* strains used in this study are described in Supplementary Table 2. Gene-deletion strains were obtained by resistance marker-guided recombination and cross-mating methods. Cells were cultured at 26°C in yeast-peptone-dextrose (YPD) or synthetic dextrose minimal medium (SDM: 0.67% yeast nitrogen base, 2% dextrose and required amino acids) at 200 rpm agitation. SILAC media were supplemented with light lysine (^12^C_6_ ^14^N_2_ L-Lys) and arginine (^12^C_6_ ^14^N_4_ L-Arg), or heavy isotopes of lysine (^13^C_6_ ^15^N_2_ L-Lys) and arginine (^13^C_6_ ^15^N_4_ L-Arg). For spot plate dilution assay, cells were grown overnight and normalized to OD_600_ of 1.0 and sequentially diluted at 1:10 dilution onto SDM agar plates containing amino acid dropout mixture in the absence or presence of 0.5 μg/ml tunicamycin, 2.5 mM NaH_2_AsO_4_, and 200 mM hydroxyurea. For H_2_O_2_ sensitivity assay, yeast cells grown to mid-log phase (OD_600_ of 0.7) were diluted to OD_600_ of 0.025 with 1.5 mM H_2_O_2_ in SDM and terminal OD_600_ of cells was measured after 1-3 days of incubation. For bacterial cultures, the yeast kinase and ubiquitin were heterolologously co-expressed in *E. coli* Rosetta 2 by 1 mM IPTG induction at 26°C for 16 h.

### Protein preparation

#### Protein precipitation

Yeast proteins were precipitated by adding ice-cold 10% trichloroacetate in TE buffer (2 mM EDTA, 10 mM Tris-HCl, pH 8.0), washed with 100% acetone, and lyophilized by vacuum centrifugation. Dessicated protein was resolubilized in urea sample buffer.

#### Protein oxidation

For oxidation assay, cells resuspended in ice-cold lysis buffer (50 mM Tris-HCl pH 7.5, 150 mM NaCl, 20 mM iodoacetamide, 1X Roche EDTA-free protease inhibitor) were disrupted by glass-bead agitation. Oxidized (carbonylated) proteins in the clarified lysate were derivatized in the dark with 2 mM 2-4-dinitrophenylhydrazine (DNPH) for 30 min in the presence of 6% SDS. The reaction was stopped by neutralization with 1 M Tris, 15% glycerol and 10% β-mercaptoethanol.

#### Recombinant protein expression and purification of kinases

N-terminally tagged Vhs1 or Sks1 were produced in C41(DE3) cells cultured in LB Media. Cells were induced at an OD_600_ of 0.6 with 1 mM IPTG and allowed to express for 4 hours at 37°C. Cells were pelleted by centrifugation at 6,000 x g for 25 min and flash frozen. Prior to purification, the cells were thawed on ice, resuspended in 5 ml of lysis buffer [50 mM Tris pH 8.0, 150mM NaCl, 10 mM imidazole, 2mM βME, complete protease inhibitors (Roche, Basel, Switzerland), 1 μg/ml DNase, 1 μg/ml lysozyme, and 1 mM PMSF] per 1 g of cells, and sonicated (21 min total, 5 s on and 10 s off). Cell lysates were cleared by centrifugation (50,000 x g for 30 min at 4°C) and filtered through a 0.45 μM filter. For purification, lysates were applied to Ni-NTA resin (Thermo Scientific, Rockford, IL) that had been equilibrated with lysis buffer containing 20 mM imidazole. The protein was eluted with lysis buffer containing 400 mM imidazole. Protein was buffer exchanged to remove imidazole [50mM Tris pH 8.0, 100mM NaCl, and 2mM βME] and the purification tag was cleaved. The protein was loaded on a HiPrep Q FF Anion Exchange Column (GE Healthcare Life Sciences, Marlborough, MA) and eluted by buffer with 300 mM NaCl. Ubiquitin-containing fractions were pooled, concentrated and further purified by size exclusion chromatography using a HiLoad Superdex 75 pg column (GE Healthcare Life Sciences, Marlborough, MA). The protein was eluted in a buffer containing 50mM Tris pH 7.5, 150mM NaCl, and 2mM DTT). Protein containing fractions were pooled and concentrated by centrifugation to 1 mM.

### Western blotting

Proteins in Laemmli buffer (for *E. coli* lysates) or urea sample buffer (for TCA-precipitated yeast proteins) containing 10% β-mercaptoethanol were resolved in 12-14% Bis-Tris PAGE gel by electrophoresis. Separated proteins were transferred onto PVDF membrane (0.2 μm, GE Healthcare Amersham) and immunoblotting was performed using the following primary antibodies: anti-ubiquitin (1:10,000; LifeSensors; MAb; clone VU-1), anti-K48 (1:10,000; Cell Signaling; RAb; clone D9D5), anti-K63 (1:4,000; EMD Millipore; RAb; clone apu3), anti-G6PDH (1:10,000; Sigma; RAb), anti-DNP (1:8,000; Sigma). Anti-mouse or anti-rabbit secondary antibodies conjugated with IRDye were purchased from LI-COR. Blots were visualized using Odyssey CLx (LI-COR Biosciences).

### Mass spectrometry

SILAC-based mass spectrometry for quantitation and mapping of ubiquitin phosphorylation sites was performed as previously described *(34, 35)*. Briefly, an equal amount of cells (labelled with either light or heavy Arg and Lys) expressing endogenous 3XFLAG-RPS31 and 3XFLAG-RPL40B were harvested from mid-log phase and disrupted by bead beating using ice-cold lysis buffer (50 mM Tris-HCl, pH 7.5, 150 mM NaCl, 5 mM EDTA, 0.2% NP-40, 10 mM iodoacetamide, 1X EDTA-free protease inhibitor cocktail (Roche), 1mM phenylmethylsulfonyl fluoride, 20 μM MG132, 1X PhosStop (Roche), 10 mM NaF, 20 mM BGP, and 2 mM Na3VO4). Lysate was clarified by centrifugation at 21,000 x *g* for 10 min at 4°C and supernatant was transferred into a new tube and diluted with threefold volume of ice-cold TBS (50 mM Tris-HCl, pH7.5, 150 mM NaCl). 3XFLAG-ubiquitin in 12-ml diluted lysate was enriched by incubation with 50 μl of EZview anti-FLAG M2 resin slurry (Sigma) for 2 h at 4°C with rotation. The resin was washed three times with cold TBS and incubated with 90 μl elution buffer (100 mM Tris-HCl, pH 8.0, 1% SDS) at 98°C for 5 min. Collected eluate was reduced with 10mM DTT, alkylated with 20 mM iodoacetamide, and precipitated with 300 μl PPT solution (50% acetone, 49.9% ethanol, and 0.1% acetic acid). Light and heavy protein pellets were dissolved with Urea-Tris solution (8 M urea, 50 mM Tris-HCl, pH 8.0). Heavy and light samples were combined, diluted four fold with water, and digested with 1 μg MS-grade trypsin (Gold, Promega) by overnight incubation at 37°C. Phosphopeptides were enriched by immobilized metal affinity chromatography (IMAC) using Fe(III)-nitrilotriacetic acid resin as previously described *(36)* and dissolved in 0.1% trifluoroacetic acid. The phosphopeptide solutions were loaded on a capillary reverse phase analytical column (360 μm O.D. x 100 μm I.D.) using a Dionex Ultimate 3000 nanoLC and autosampler and analyzed using a Q Exactive Plus mass spectrometer (Thermo Scientific). Data collected were searched using MaxQuant (ver. 1.6.5.0) and chromatograms were visualized using Skyline (ver. 20.1.0.31, MacCoss Lab).

### *In vitro* kinase activity

Kinase activity assays were performed in a reaction mixture containing 50 mM Tris (pH 7.4), 150 mM NaCl, 10 mM MgCl_2_, 0.1 mM ATP, 1 mM DTT, 0.5 μM ubiquitin, and 50 nM kinase. The reaction mixture was carried out at 30°C for 30 min and quenched by adding equal volume of Laemmli buffer with 10% β-mercaptoethanol and heating at 98°C for 10 min.

## ACKNOWLEDGEMENTS

We are very grateful to K. Rose for technical advice and assistance with quantitative mass spectrometry analysis. We are also grateful to B. Brasher technical advice and recommending reagents. We also thank T. Graham for critical reading of the manuscript. J.M.T. was funded by NIH training grant T32 CA119925. This research was supported by NIH grant R21 AG053562 (to J.A.M.), R01 GM118491 (to J.A.M.), and R35 GM118089 (to W.J.C.).

---

**Figure 1, supplement 1.**
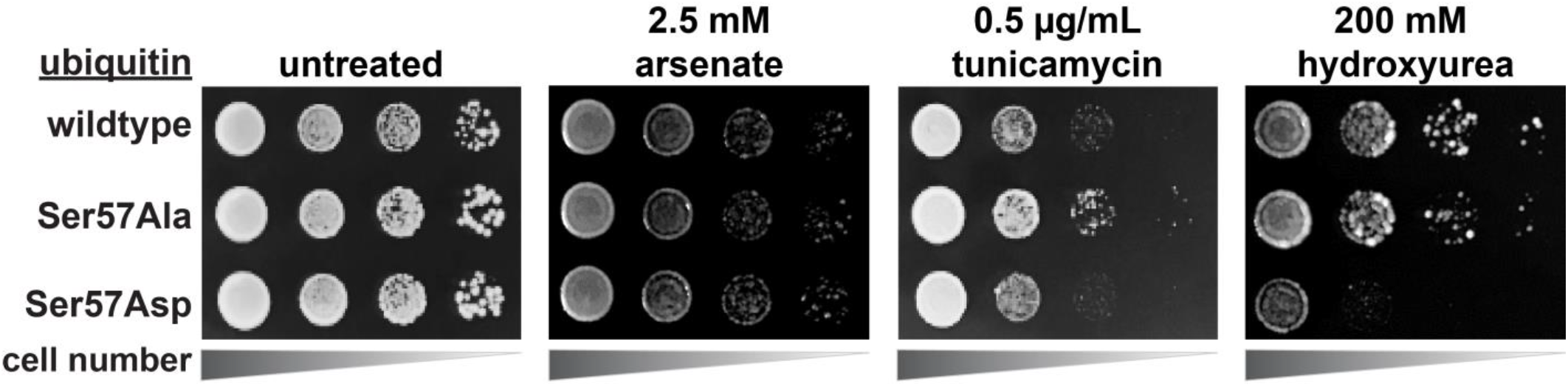
Phenotypic analysis of yeast cells expressing exclusively WT, S57A, or S57D ubiquitin. Yeast cells (strain background SUB280) were grown overnight and normalized to OD_600_ of 1.0 and sequentially diluted at 1:10 dilution onto synthetic dextrose medium (SDM) agar plates containing amino acid dropout mixture in the absence or presence of 2.5 mM NaH2AsO4, 0.5 μg/ml tunicamycin, and 200 mM hydroxyurea.

**Figure 1, supplement 2.**
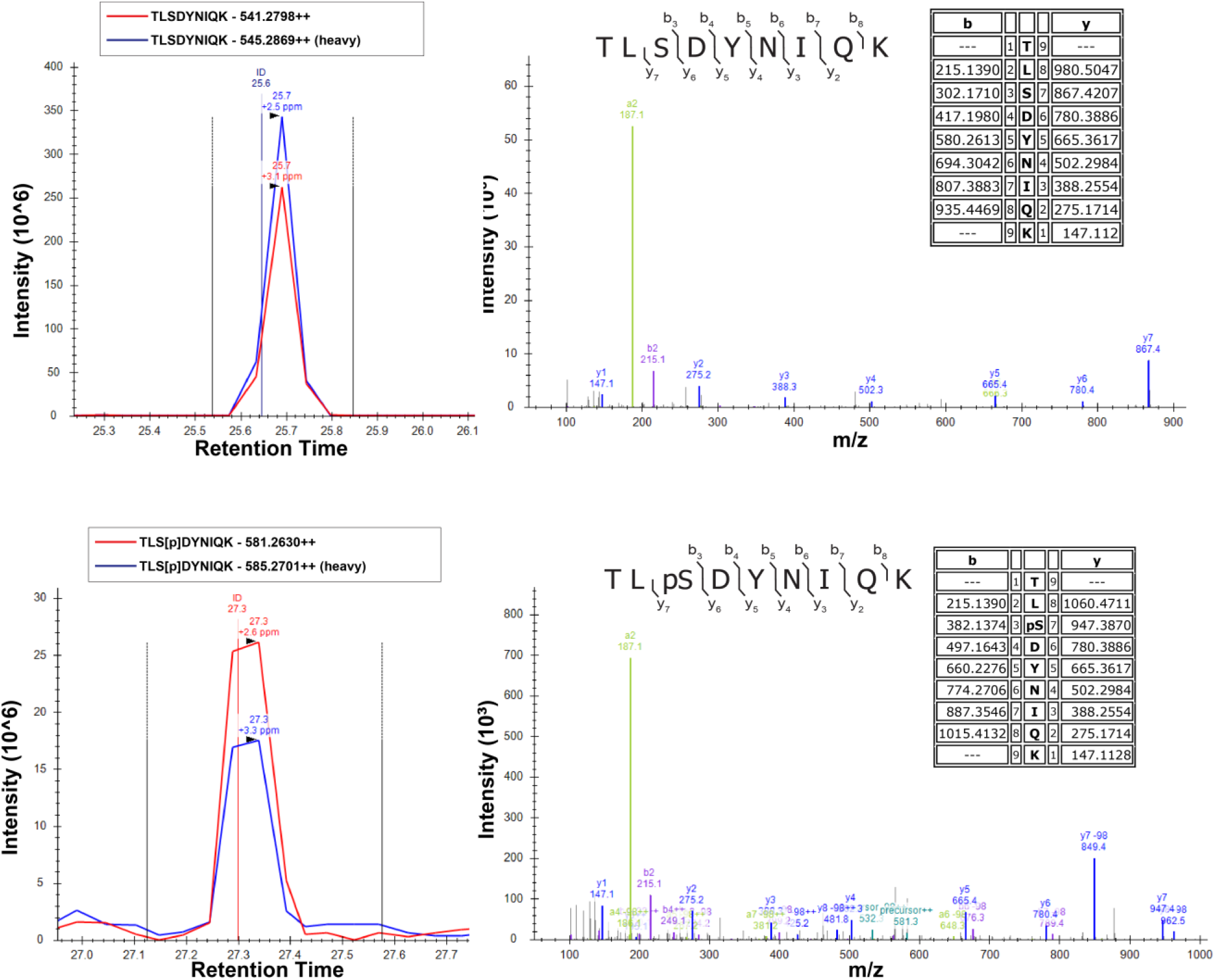
SILAC-MS fragmentation spectra of unmodified (top) and Ser57-phosphorylated (bottom) peptides of ubiquitin isolated from yeast cells grown in the presence (light) or absence (heavy) of H_2_O_2_. Cells were grown in SILAC media (supplemented with light or heavy lysine and arginine) to mid-log phase (OD_600_ of 0.6-0.7) and treated with 0.6 mM H_2_O_2_ for 30 minutes. Chromosomally expressed 3xFLAG-tagged ubiquitin (from the *RPS31* and *RPL40B* loci) was isolated by affinity purification and digested with trypsin. Phosphopeptides were enriched by immobilized metal affinity chromatography (IMAC), separated by a capillary reverse phase analytical column, and analyzed on a Q Exactive mass spectrometer. Theoretical masses of *b/y* ions are tabulated and MS-observed masses are shown in the spectra.

**Figure 2, supplement 1.**
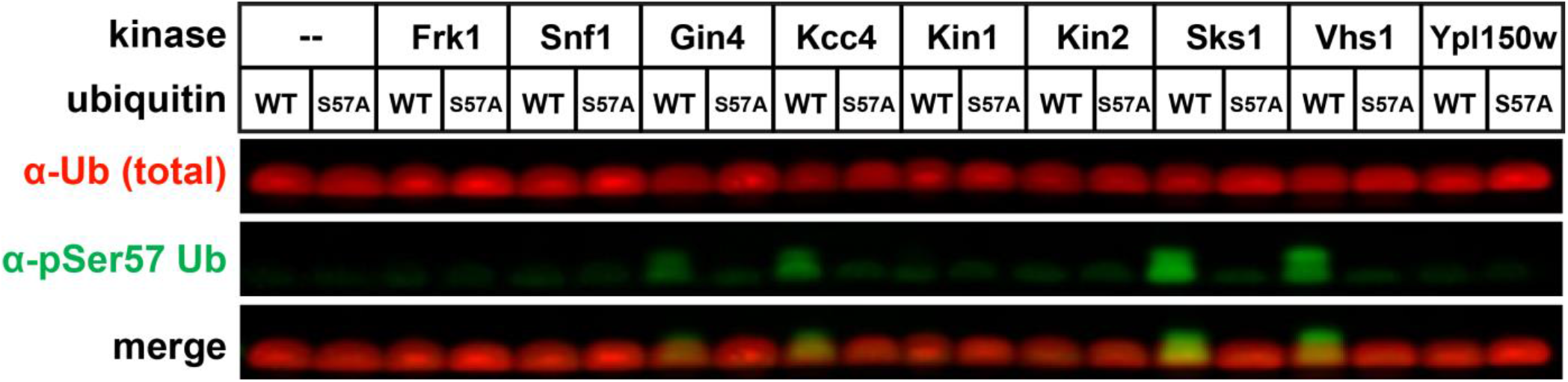
Anti-phospho-Ser57 western blot of *E. coli* (Rosetta 2) whole-cell lysates after heterologous co-expression of ubiquitin and yeast members of the Snf1-related family of kinases.

**Figure 2, supplement 2.**
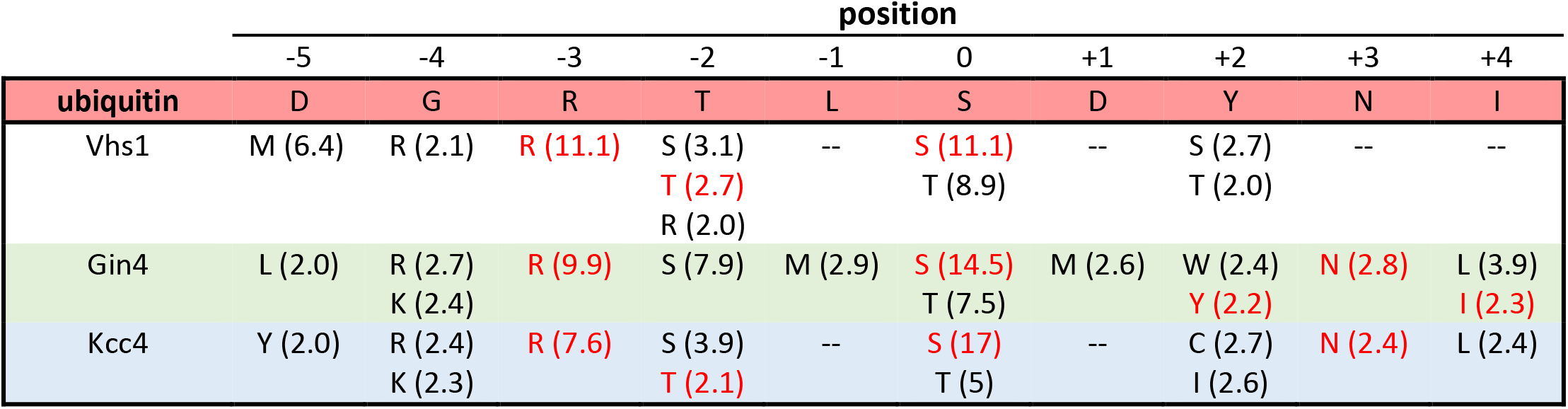
For yeast Ser57 ubiquitin kinases, we analyzed consensus phosphorylation motifs as determined from a previous study based on in vitro activity analysis on peptide libraries [33]. Values in parentheses are the quantified selectivity values, based on site preference of in vitro activity. Only amino acids selected at a position with a value >2 are shown.

**Figure 2, supplement 3.**
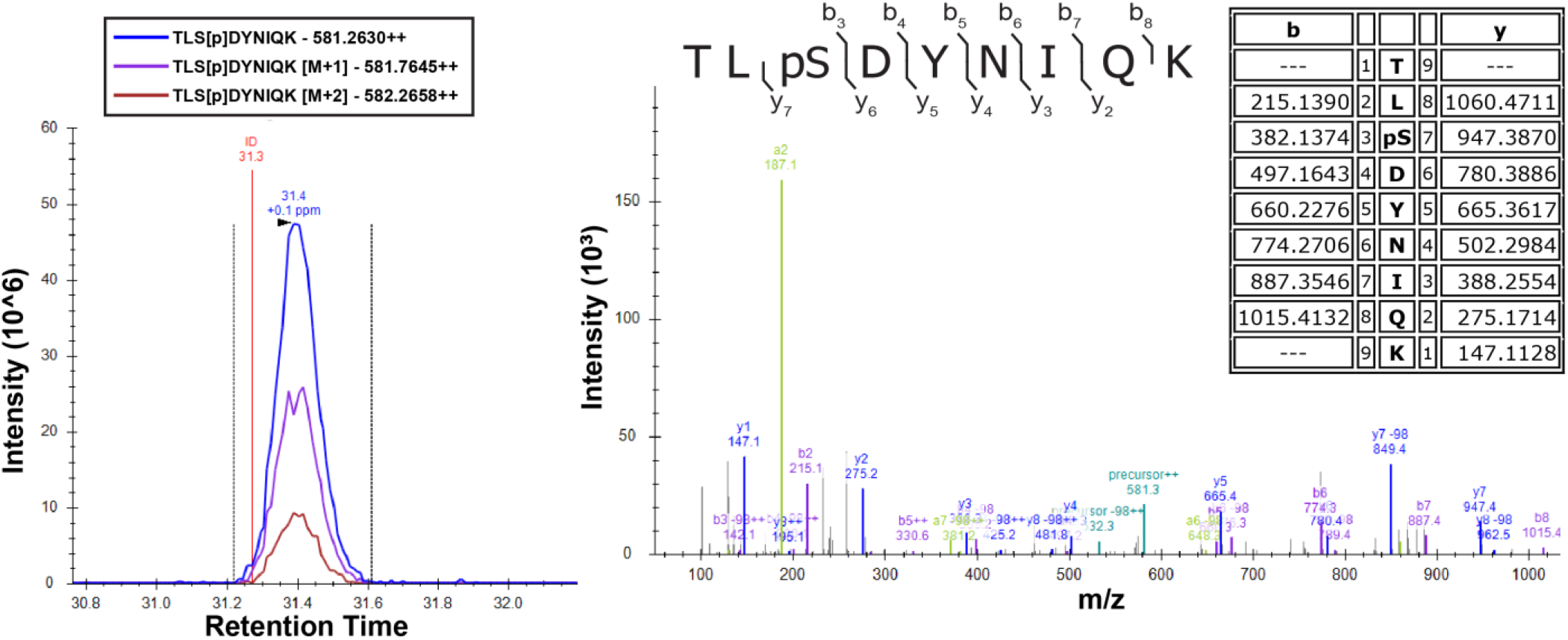
Detection of Ser57 phosphorylation of ubiquitin in the presence of Vhs1 by mass spectrometry. Vhs1 and ubiquitin of yeast were co-expressed in *E. coli* Rosetta 2 by IPTG induction. Ubiquitin was isolated using UBD-capture pull-down, digested with trypsin, and analyzed on a Q Exactive mass spectrometer. Theoretical masses of *b/y* ions are tabulated and MS-observed masses are shown in the spectra.

**Figure 2, supplement 4.**
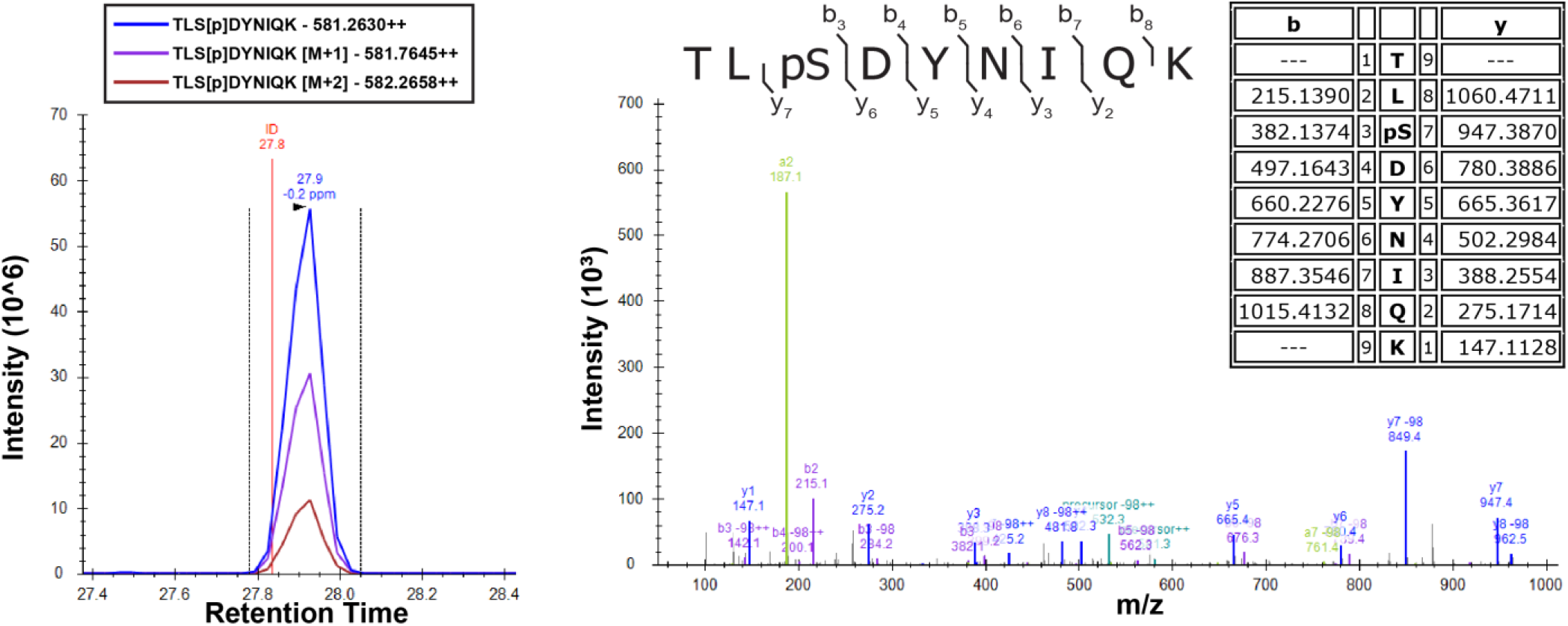
Detection of Ser57 phosphorylation of ubiquitin in the presence of Sks1 by mass spectrometry. *In vitro* kinase activity assay was carried out using purified recombinant Sks1 and tetra-ubiquitin as described in the *Materials and Methods.* Total peptide digests of the reaction mixture were separated by reverse-phase capillary column and analyzed on a Q Exactive mass spectrometer. Theoretical masses of *b/y* ions are tabulated and MS-observed masses are shown in the spectra.

**Figure 2, supplement 5.**
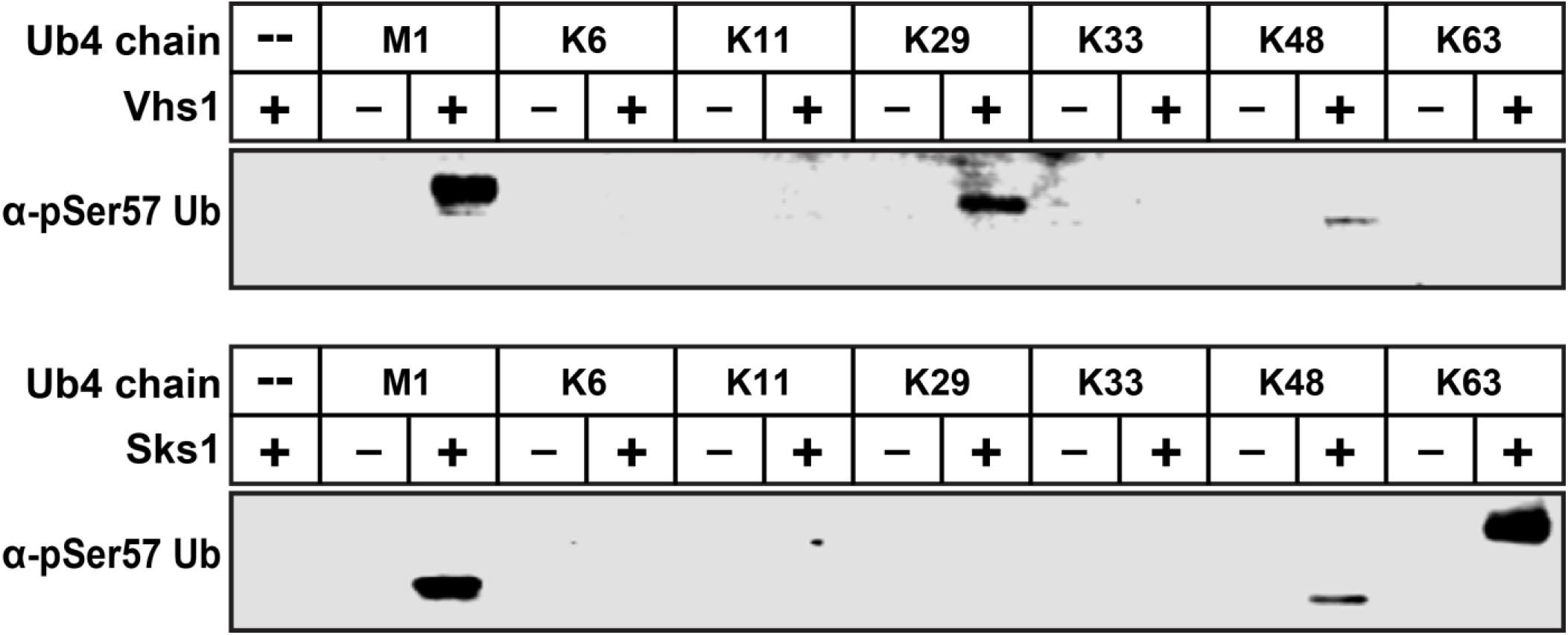
Vhs1 and Sks1 have distinct linkage type preferences. *In vitro* reconstitution assay was carried out by mixing recombinant Vhs1 (D) or Sks1 (E) and tetra-ubiquitin with different linkage types (M1, K6, K11, K29, K33, K48, K63) as described in the *Materials and Methods.* Ser57 phosphorylation was detected by western blot using anti-phospho-Ser57 ubiquitin antibody.

**Figure 2, supplement 6.**
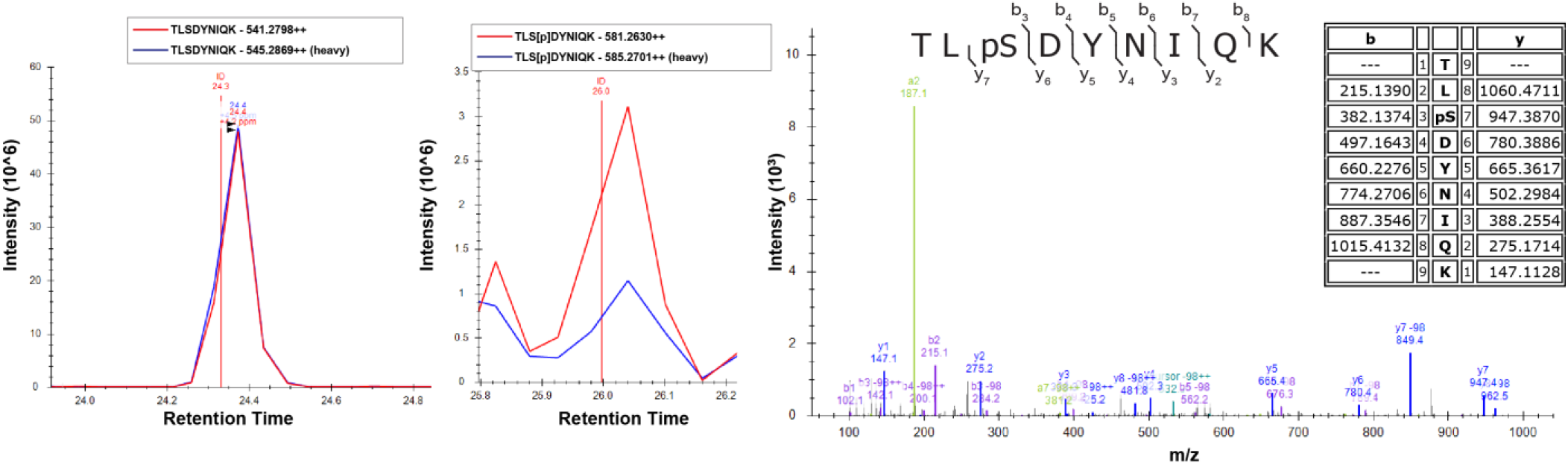
Overexpression of Sks1 in yeast cells increases Ser57 phosphorylation of ubiquitin. Cells were grown in SILAC media supplemented with light or heavy lysine and arginine. Chromosomally expressed 3xFLAG-tagged ubiquitin (from the *RPS31* and *RPL40B* loci) was isolated by affinity purification and digested with trypsin. Phosphopeptides were enriched by immobilized metal affinity chromatography (IMAC), separated by a capillary reverse phase analytical column, and analyzed on a Q Exactive mass spectrometer. Theoretical masses of *b/y* ions are tabulated and MS-observed masses are shown in the spectra.

**Figure 3, supplement 1.**
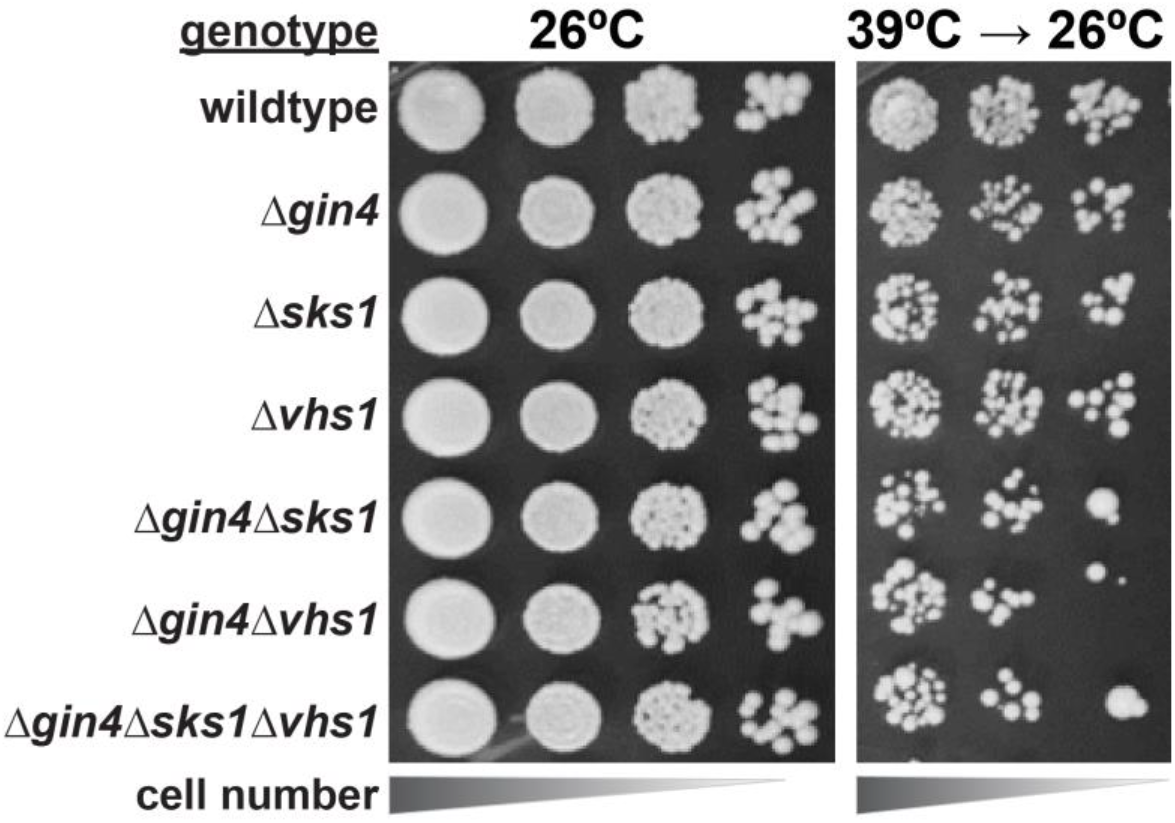
Analysis of cell tolerance to heat stress in yeast kinase deletion cells. Null-mutants derived from SEY6210 yeast strain were serially diluted onto YPD plates and incubated at 39°C for 18 h and shifted to 26°C for growth recovery.

**Figure 3, supplement 2.**
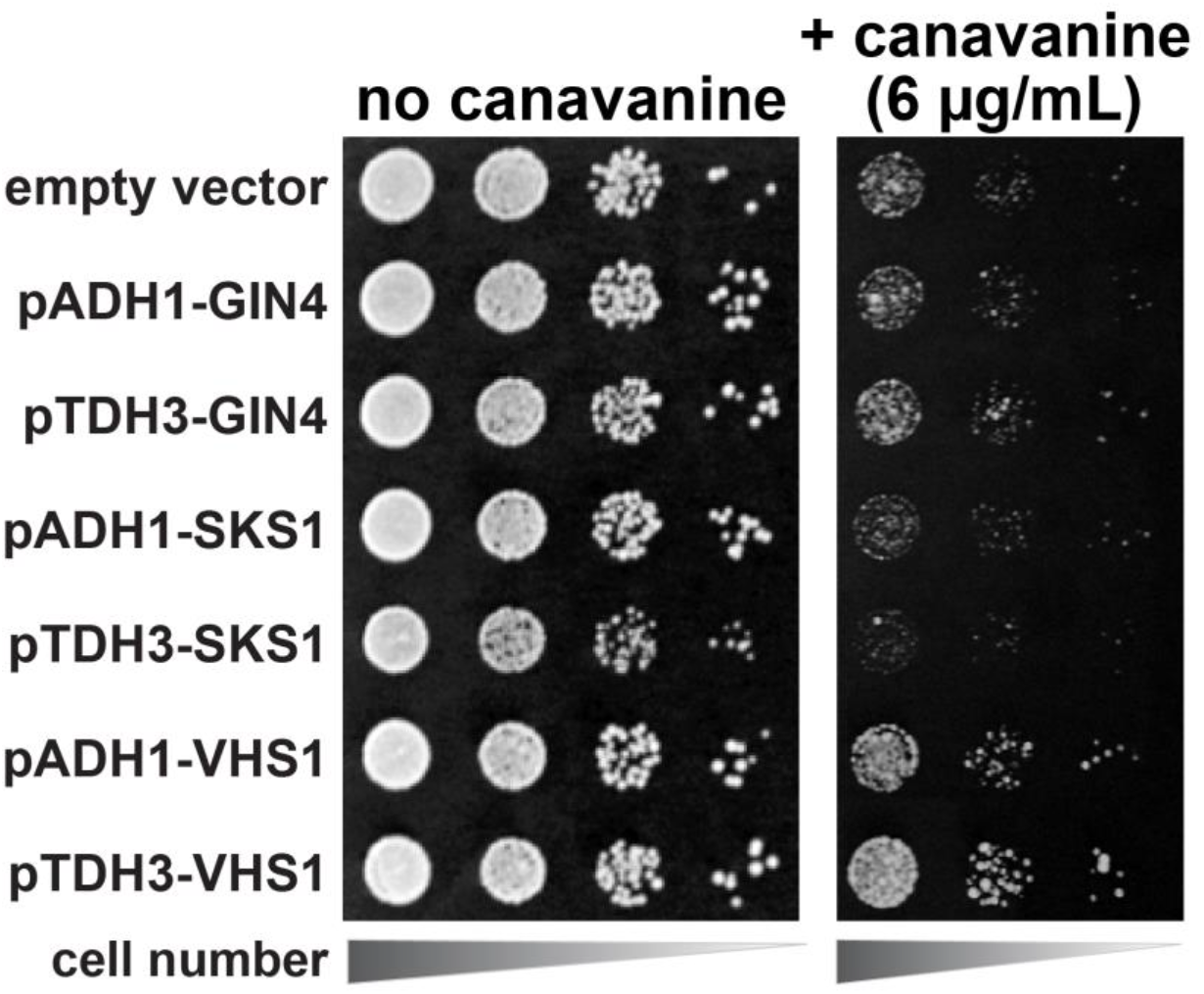
Overexpression of Vhs1 slightly increases yeast tolerance to canavanine. SEY6210 cells were transformed with Gin4, Sks1, or Vhs1, controlled by *pADH1* or *pTDH3* constitutive promoter. Transformants were serially diluted onto dropout SDM in the presence or absence of 6 μg/ml canavanine.

**Figure 3, supplement 3.**
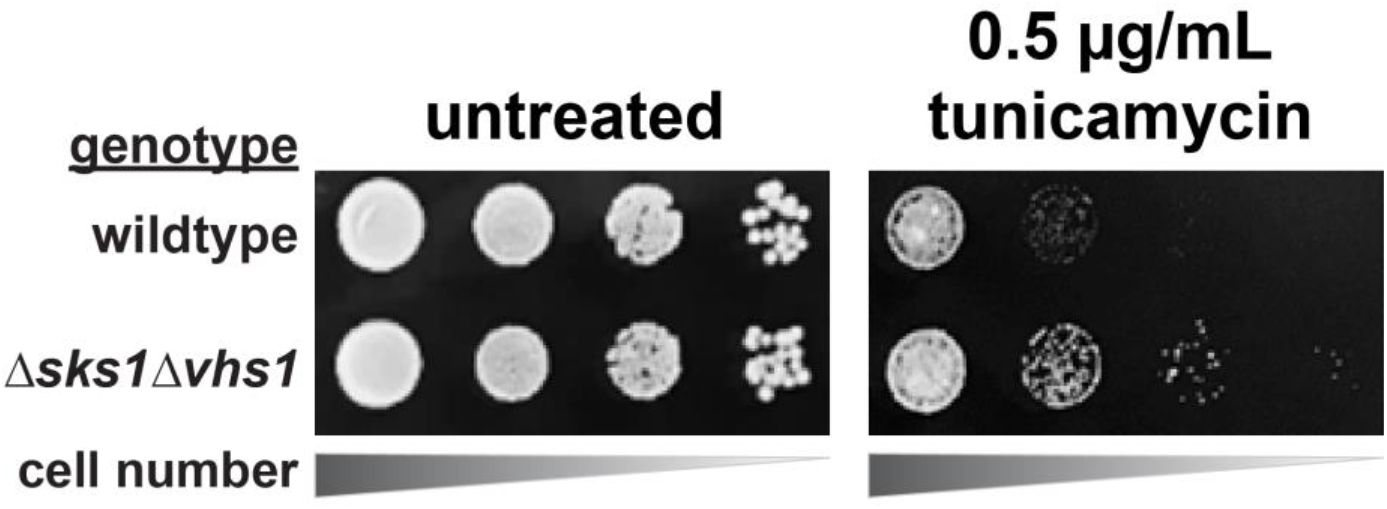
*SKS1* and *VHS1* regulate the growth response of yeast to tunicamycin. SEY6210 (wt) and *Δsks1Δvhs1* mutant strains were serially diluted onto YPD with or without 0.5 μg/ml tunicamycin supplementation.

**Figure 3, supplement 4.**
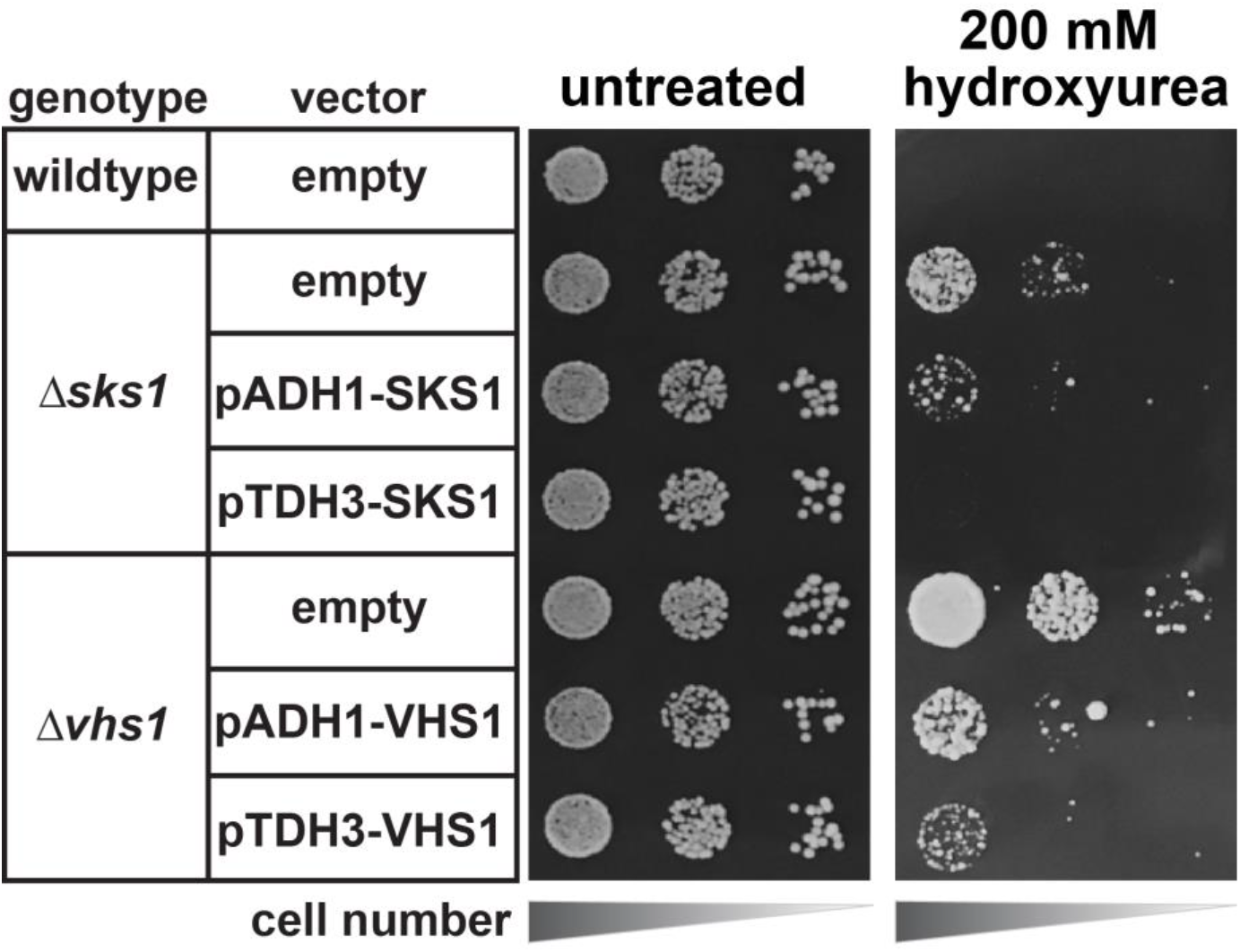
Sks1 and Vhs1 expression determines yeast sensitivity to hydroxyurea. Mutant cells derived from SEY6210 were serially diluted onto dropout SDM agar plate in the presence or absence of 0.2 M hydroxyurea.

**Figure 4, supplement 1.**
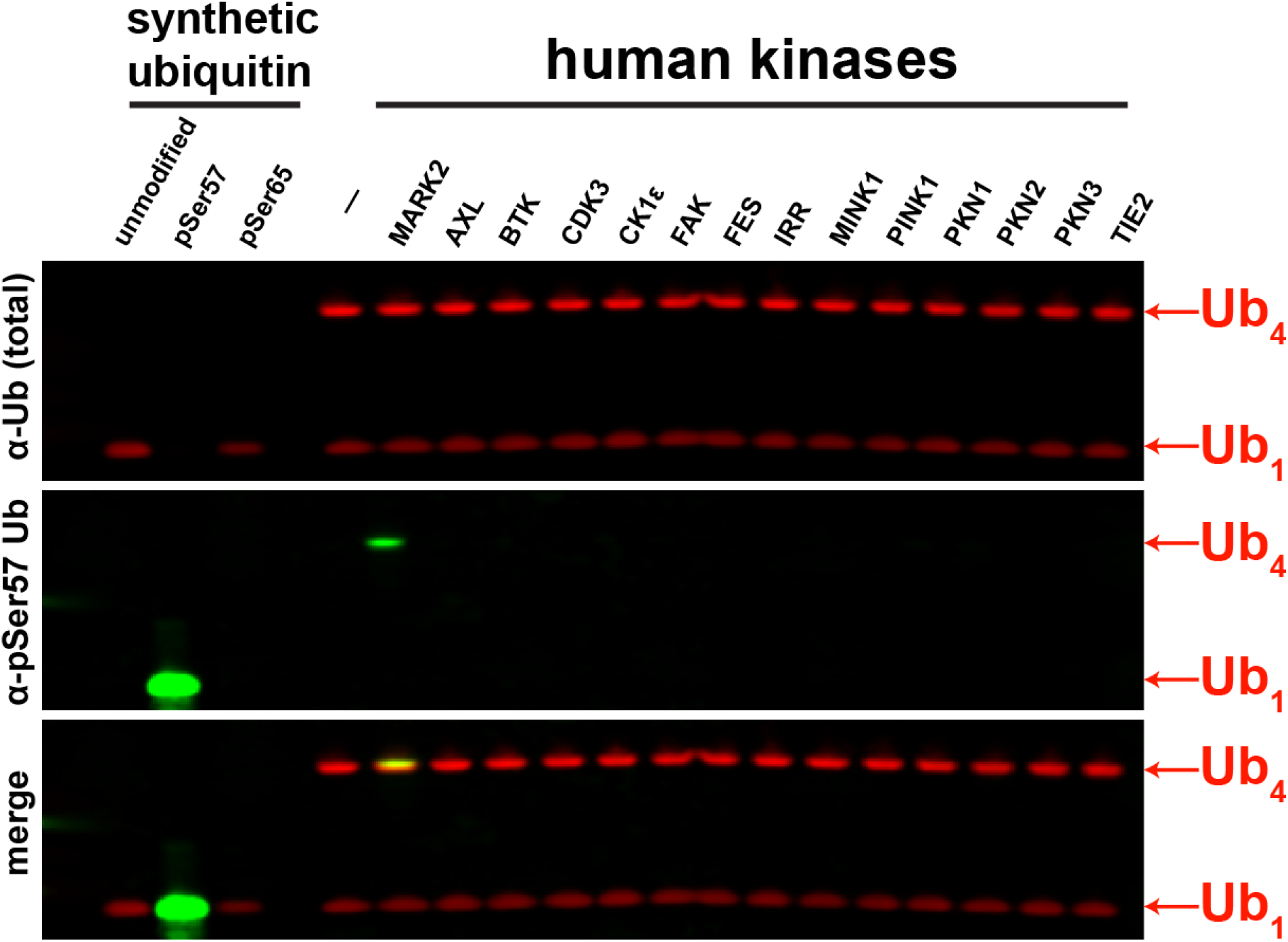
Screening of human ubiquitin Ser57 kinases by Western Blot. Select kinases of different superfamilies were used for *in vitro* kinase activity assays using tetra-ubiquitin. Ubiquitin phosphorylation at the Ser57 residue was determined by Western blot using anti-phospho-Ser57 ubquitin antibody.?

**Figure 4, supplement 2.**
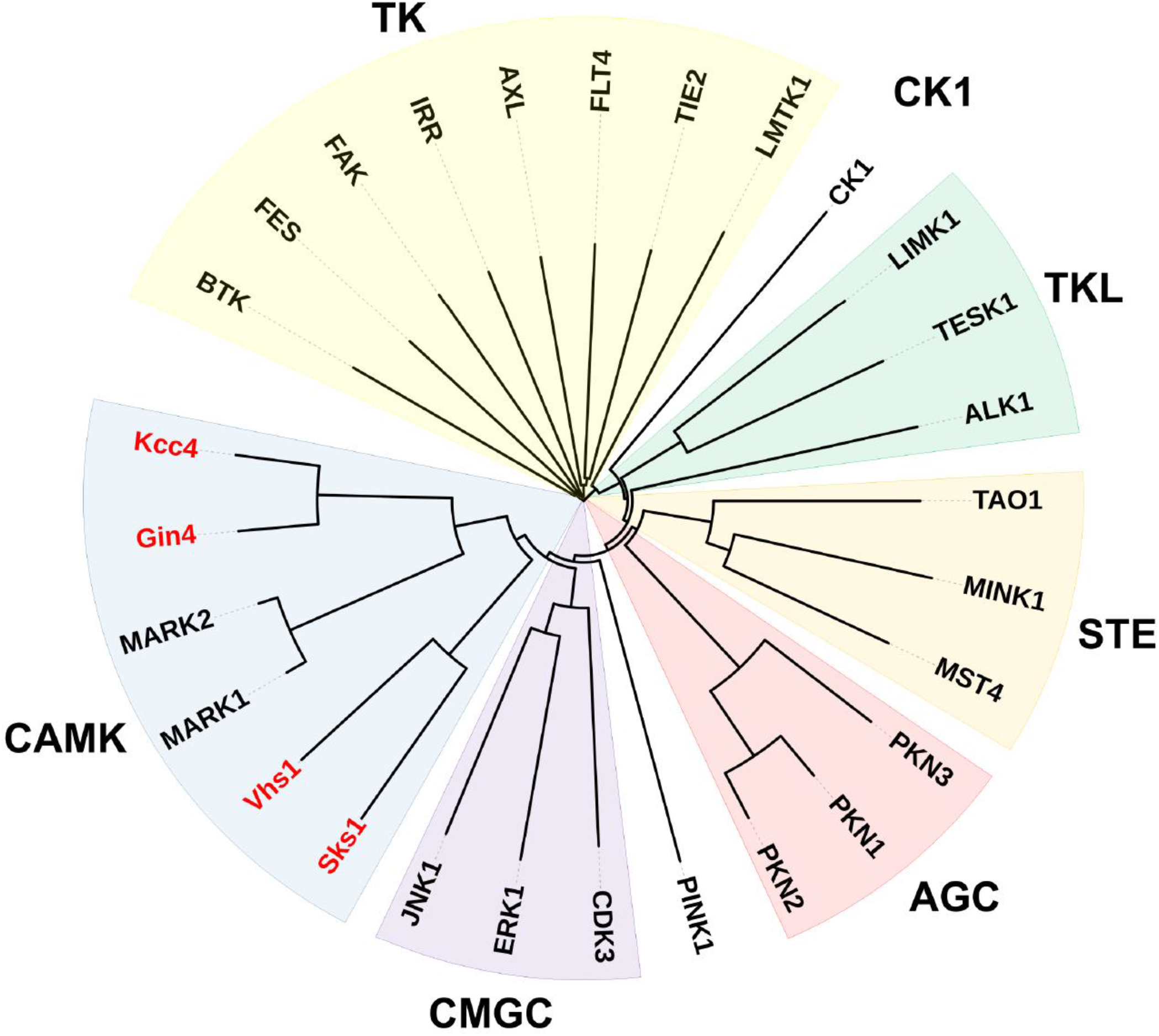
Yeast ubiquitin kinases Sks1, Vhs1, Gin4 and Kcc4 cluster with human MARKs of the CAMK superfamily of kinases. Sequence alignment and phylogenetic neighboring of kinase domains were analyzed by Clustal Omega (EMBL-EBI) and the circular rooted phylogram was constructed using iTOL. Yeast kinases are shown in red.

